# Glioma grade map: a machine-learning based imaging biomarker for tumor characterization

**DOI:** 10.1101/133249

**Authors:** András Jakab, Péter Molnár, Miklós Emri, Ervin Berényi

## Abstract

**Purpose:** To use T1-, T2-weighted and diffusion tensor MR images to portray glioma grade by employing a voxel-wise supervised machine learning approach, and to assess the feasibility of this tool in preoperative tumor characterization.

**Materials and Methods:** Conventional MRI, DTI datasets and histopathological evaluations of 40 patients with WHO grade II-IV gliomas were retrospectively analyzed. Databases were construed incorporating preoperative images, tumor delineation and grades. This data was used to train a multilayer perceptron based artificial neural network that performed voxel-by-voxel correlation of tumor grade and the feature vector. Results were mapped to grayscale images, whereas grade map was defined as a composite image that depicts grade assignments for intra-tumoral regions. The voxel-wise probability for high grade tumor classification was calculated for the entire tumor volumes, defined as the grade index.

**Results:** The color hue on glioma grade maps allowed the discrimination of low and high grade cases. This method revealed connection between the heterogeneous appearance of tumors and the histopathological findings. Classification by the grade index had 92.31% specificity, 85.71% sensitivity.

**Conclusion:** Glioma grade maps are advantageous in the visualization of the heterogeneous nature of intra-tumoral diffusion and relaxivity and can further enhance the characterization of tumors by providing a preoperative modality that expands information available for clinicians.

**ABBREVIATIONS:**

ADC: apparent diffusion coefficient;
ANN: artificial neural networks;
DWI: diffusion weighted imaging;
DTI: diffusion tensor imaging;
FA: fractional anisotropy;
HGPM: high grade probability map;
LGPM: low grade probability map;
TPM: tumor probability map;
WHO: World Health Organization

## INTRODUCTION

The most prevalent forms of brain tumors are glial neoplasms, whereas astrocytic tumors constitute the majority of gliomas, as stated by the World Health Organization (WHO) classification (Cohen and Weller, 2007); however, mixed cellular composition is also common (Ohgaki et al., 2005). Separating gliomas into low-grade and high-grade classes has become the means for assessing the neoplastic biological behavior, and this partitioning fundamentally determines therapy and patients’ survival.

Magnetic resonance imaging (MRI) bears remarkable tissue contrast and depicts morphological details of brain malignancies. Despite the endeavors to use conventional MRI for the delineation of high-grade gliomas, low sensitivity and specificity was reported (Law et al., 2003). Diffusion-weighted imaging probes the motion of molecules by applying diffusion sensitizing magnetic gradients (Le Bihan et al., 1986) while diffusion tensor imaging (DTI) allows estimating the direction of the diffusion and the degree of anisotropy. Such measures are often correlated with biological features of brain tumors; this is hallmarked by numerous observations made on the relationship between DTI derived values and histopathological findings. The rationale of such efforts if to quantify the microscopic properties of diffusion and to compare these values with tissue properties like tumor cellularity, cell density, proliferation activity (Sugahara et al., 1999; Beppu et al., 2005, Sadeghi et al., 2008) or differentiate astrocytomas from oligodendrogliomas (Khayal et al., 2011). Diffusion tensor imaging derived parameters were extensively tested for clinical usability in determining the grade of CNS gliomas (i.e. low or high; or anaplastic vs. low-grade), this is mainly achieved by collecting quantitative parameters from preoperative images (Inoue et al. 2005; Higano et al., 2006; Goebell et al., 2006; Tozer et al., 2007; Jakab et al., 2011; Ellingson et al., 2011; Haegler et al., 2011). Successful classification was achieved when using a combination of diffusion parameters (multiparametric approach) or by more complex dissection of imaging data (Murakami et al., 2009; Emblem et al., 2008; Raab et al., 2010).

It is possible to visualize biologically diverse regions within a tumor based on image analysis and various modeling approaches; a method was reported that depicts histological subtypes (i.e. “oligo-like” or “astro-like” regions, according to the authors' nomenclature) of low-grade gliomas as color maps (Khayal et al., 2009). Similarly, “nosologic images” graphically represent different tumor types by performing complex interpretation of MR spectroscopy data (De Edelenyi et al., 2000; Luts et al., 2009) and it was practical to use T2 and ADC values for tumor xenograft characterization by segmenting tumor images into various sub-populations (Carano et al., 2004).

We hypothesize that machine-learning algorithms are capable of integrating information from preoperative images whilst multidimensional pattern-recognition techniques could enhance the characterization of gliomas. A practical approach is supervised machine learning where previously determined ground truth is provided by histopathology, and mathematical models are optimized for finding the correlation of individual, subject-based data and the tumor classification. One such method, the artificial neural networks (ANN), has long been investigated as a potential candidate for oncology decision support (Baxt, 1995; Hagberg, 1998) finding more specific aims as brain tumor classification (Wang et al., 2006; Joshi et al., 2010). By the same token, our investigation was designed to introduce a new visualization method that portrays glioma grade by incorporating information from postgadolinium T1- and T2-weighted, diffusion-weighted and parametric images that were computed from diffusion-tensor measurements. We focused on the development of an imaging biomarker that estimates tumor grade by employing a voxel-wise computational approach based on a supervised learning algorithm.

## MATERIALS AND METHODS

The study cohort consisted of 40 consecutively recruited patients meeting the following inclusion criteria: first occurrence of intracerebral glioma (WHO grade II-IV), availability of preoperative diffusion tensor and conventional MR imaging. Patients were recruited between 2007 and 2010. No patients underwent treatment for their brain malignancies prior to radiologic examinations. Histopathological diagnoses were assigned according to the WHO classification criteria by a neuropathologist with 30 years’ experience (P.M.). Tissue samples were obtained by either stereotactic biopsy or surgical debulking. Subject characteristic data are detailed in Table 1.

**Table 1.**
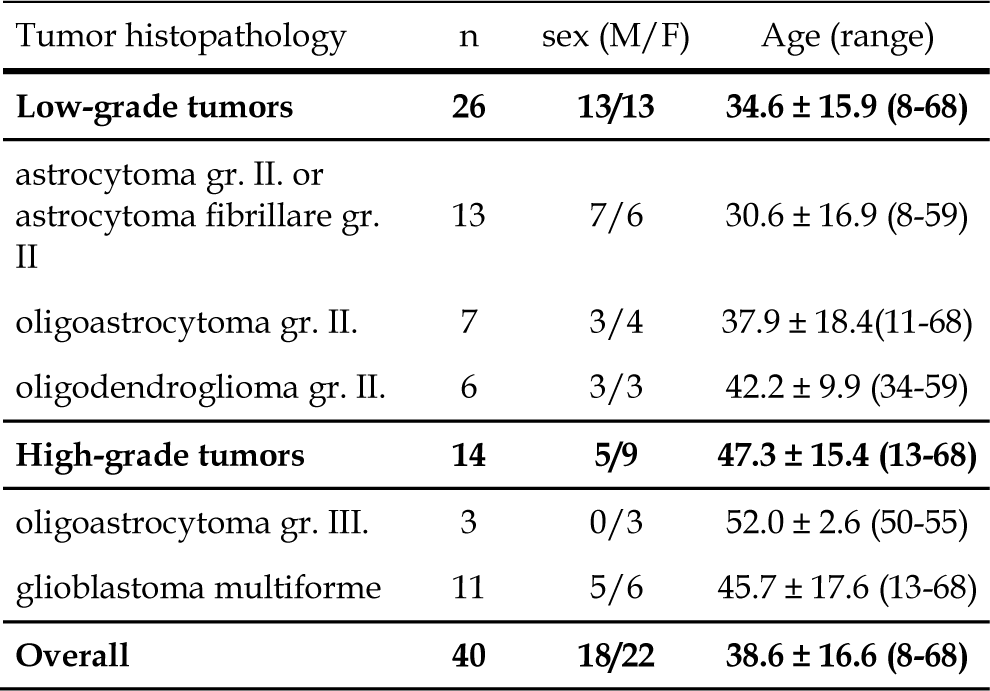
Patient baseline characteristics.

| Tumor histopathology | n | sex (M/F) | Age (range) |
| --- | --- | --- | --- |
| **Low-grade tumors** | **26** | **13/13** | **34.6 ± 15.9 (8-68)** |
| astrocytoma gr. II. or<br>astrocytoma fibrillare gr. II | 13 | 7/6 | 30.6 ± 16.9 (8-59) |
| oligoastrocytoma gr. II. | 7 | 3/4 | 37.9 ± 18.4(11-68) |
| oligodendroglioma gr. II. | 6 | 3/3 | 42.2 ± 9.9 (34-59) |
| **High-grade tumors** | **14** | **5/9** | **47.3 ± 15.4 (13-68)** |
| oligoastrocytoma gr. III. | 3 | 0/3 | 52.0 ± 2.6 (50-55) |
| glioblastoma multiforme | 11 | 5/6 | 45.7 ± 17.6 (13-68) |
| **Overall** | **40** | **18/22** | **38.6 ± 16.6 (8-68)** |

### Imaging protocol

Imaging was performed on a GE Signa 1.5-T TwinGradient whole body scanner (GE Medical Systems, Milwaukee, WI) equipped with an 8-channel phased-array head coil. For anatomical imaging, we obtained postgadolinium 3DT1 scans. The DTI dataset was acquired by using a diffusion-weighted echo-planar imaging sequence with 25 gradient directions. Acquisition parameters are summarized in Table 2.

**Table 2.** Image acquisition parameters

|  | Anatomic imaging | Diffusion tensor imaging |
| --- | --- | --- |
| Sequence | 3DT1 SPGR | SS EPI, 25 directions |
| TR / TE | 30/7 ms | 1000/98 ms |
| Voxel size | 0.625 x 0.625 mm | 1 x 1 mm |
| Slice thickness | 1.1 mm | 3.3 mm |
| Matrix | 256 x 256 | 128 x 128 |
| Field of view | 350 mm | 260 mm |

### Image processing

DTI data were used to calculate the following 6 scalar maps: T2-weighted images (DWI without diffusion sensitization – B_0_), directionally averaged raw DWI images, fractional anisotropy, longitudinal and radial diffusivity images, apparent diffusion coefficient maps. To obtain anatomical correspondence through the imaging modalities, postcontrast T1 scans were coregistered with the B_0_ images and eventually, all images were re-sampled to smaller matrices of 128 * 128 voxels. Intensity normalization of the 3DT1 images was performed with the built-in “enhance contrast” command in the ImageJ software tool (Abramoff et al., 2004). Tumor outlines were defined as the maximal abnormal region as seen on postcontrast T1 images and were validated by a neuroradiologist; manual delineation resulted in a binary tumor-mask file with dimensions identical to the resampled radiologic images. For high grade tumors, the maximal abnormal region was defined as T1 hyperintensity while the extension of low grade tumors was derived from the parallel observation of T1 and T2 images; T1 hypointensity inside the encircling T2 hyperintensity was delineated (to exclude the putative regions of oedema). Tumor segmentation, diffusion tensor calculation and coregistrations were executed using the Slicer 3D software package (Pieper et al., 2004), DTI scalar calculations were based on the equations described elsewhere (Basser and Pierpaoli, 1993).

### Data processing

We utilized image information of the 40 patients to generate two different databases for the classifier training procedure. In each database, samples represented consecutive voxels’ values on the images and a categorical variable was also assigned voxel-wise resulting in a total number of 8 variables per voxel. Database “A” provided ground truth for separating the voxels sampled from a low grade tumor or a high grade tumor. In contrast, the aim of database “B” was to separate tumorous regions from non-tumorous regions, as later described, this was only considered important for visualizing the results. Database “A” was built by sampling exclusively the intra-tumoral regions, the categorical variable was the tumor grade as determined by the histopathology workup (low grade=0; high grade=1) and this was assigned case-wise, without spatial control of the histology sampling. Database “B” included every intracerebral voxels, whereas the eighth, dichotomous variable described whether the voxel was intra-tumoral (value: 1) or of normal-appearing brain tissue (value: 0) as determined by the tumor-mask. The relationship between imaging data and the categorical variables (i.e. tumor grade, tumor or normal-appearing brain tissue) was analyzed voxel-wise by utilizing a feed-forward, back propagation multilayer perceptron artificial neural network algorithm in the SPSS 17.0 for Windows software (SPSS Inc., Chicago, IL, USA). The training regime was based on the random splitting of the dataset into three groups: training (70% of all voxels), interactive testing (20%) and independent evaluation sample for reporting the classifier accuracy, “holdout” samples (10%). This supervised learning method resulted in two distinctive models, the first aiming to predict the grade of the glioma while the other assesses if the voxel is sampled from a tumor or from the normal-appearing brain tissue.

### Grade maps

Data acquisition and image processing steps are summarized in Figure 1.

**Figure 1.**
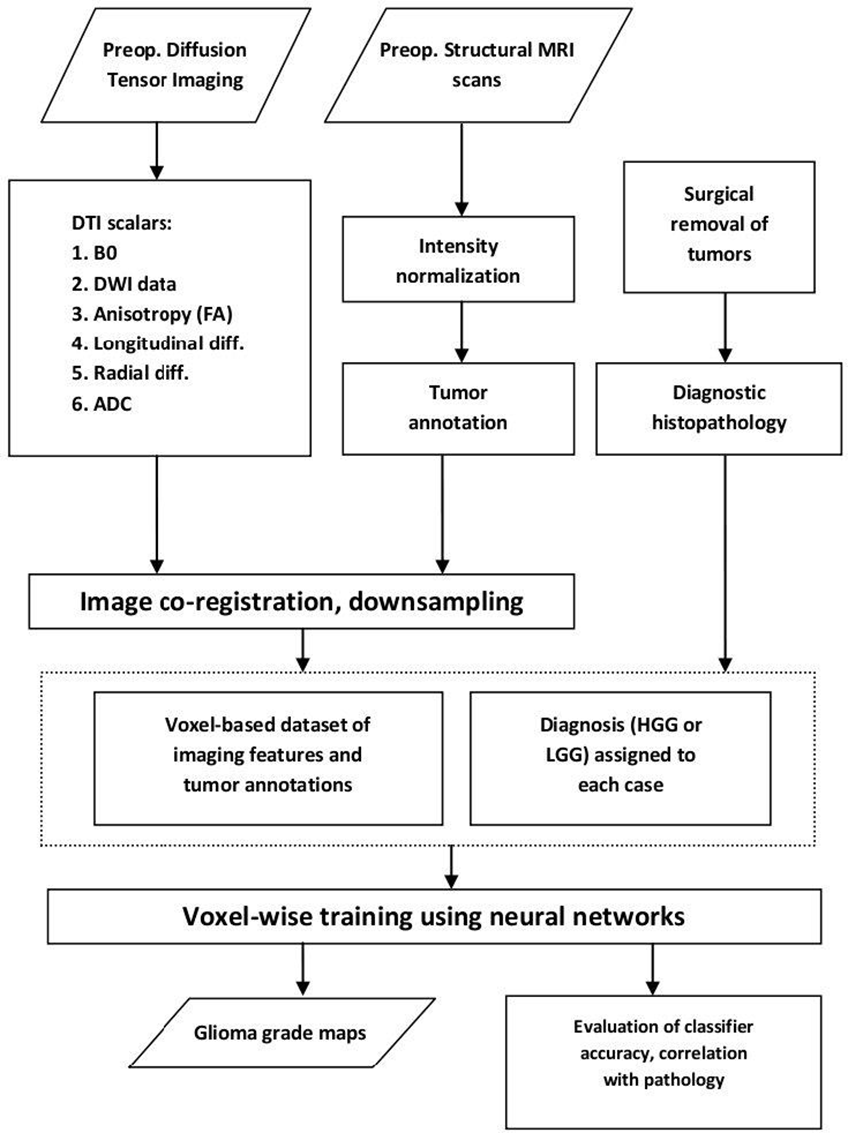
Glioma grade map generation. Work flow of the main image and data acquisition, image processing steps, numerical database creation.

Data transfer between the statistical software package and the image processing application was carried out by a using a custom short program code (i.e. macro) in ImageJ. After the classifier training, the image dataset was re-evaluated for each patient and outputs were mapped to grayscale images. The voxel-wise outputs of the neural network were continuous variables that estimated the likelihood of voxel group memberships. Grade map generation consisted of the following steps. First, we run the *a priori* trained neural net based on database “A” to generate an image yielding low- and high-grade voxel membership probability maps (LGPM and HGPM). Then, the second neural network estimation – previously trained with database “B” – resulted in an image that quantified the probability of tumor-like regions (tumor probability map, TPM). To provide a graphical representation, LGPM and HGPM images were weighted with the tumor probability maps. Eventually, we defined the glioma grade map as a color-coded composite image where the color lookup table was specified as follows. Blue shade represents low-grade regions (LGPM), red shade is for high-grade regions (HGPM), and opacity is derived from the TPM, overlaid on the co-registered anatomical T1-weighted image. The database structure for the classifier training is illustrated in Figure 2., while the intermediate gray-scale and the resulting color-coded images are exemplified in Figure 3.

**Figure 2.**
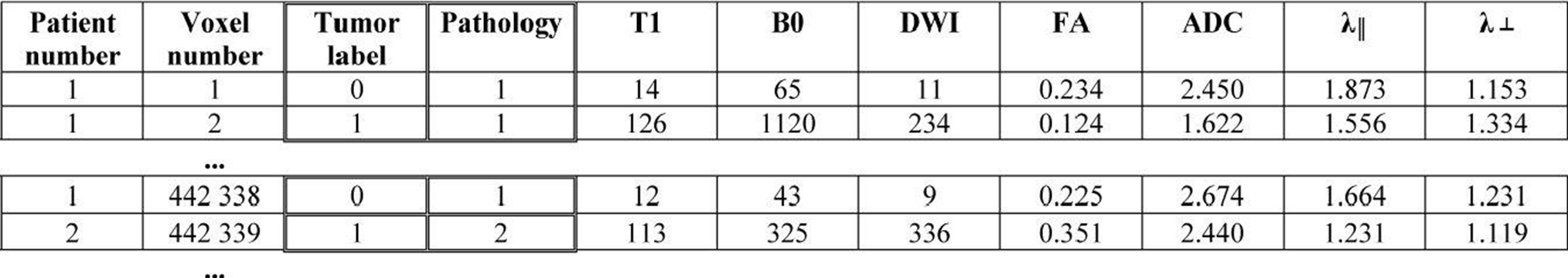
Dataset structure for training an artificial neural network classifier. Individual samples are image voxels of 40 subjects, each given a categorical variable: tumor label (e.g. 1 if the voxel was sampled from inside, or 0 if outside a tumor), histopathological diagnosis (1: low grade glioma, 2: high grade glioma). Values of 6 imaging features are exemplified.

**Figure 3.**
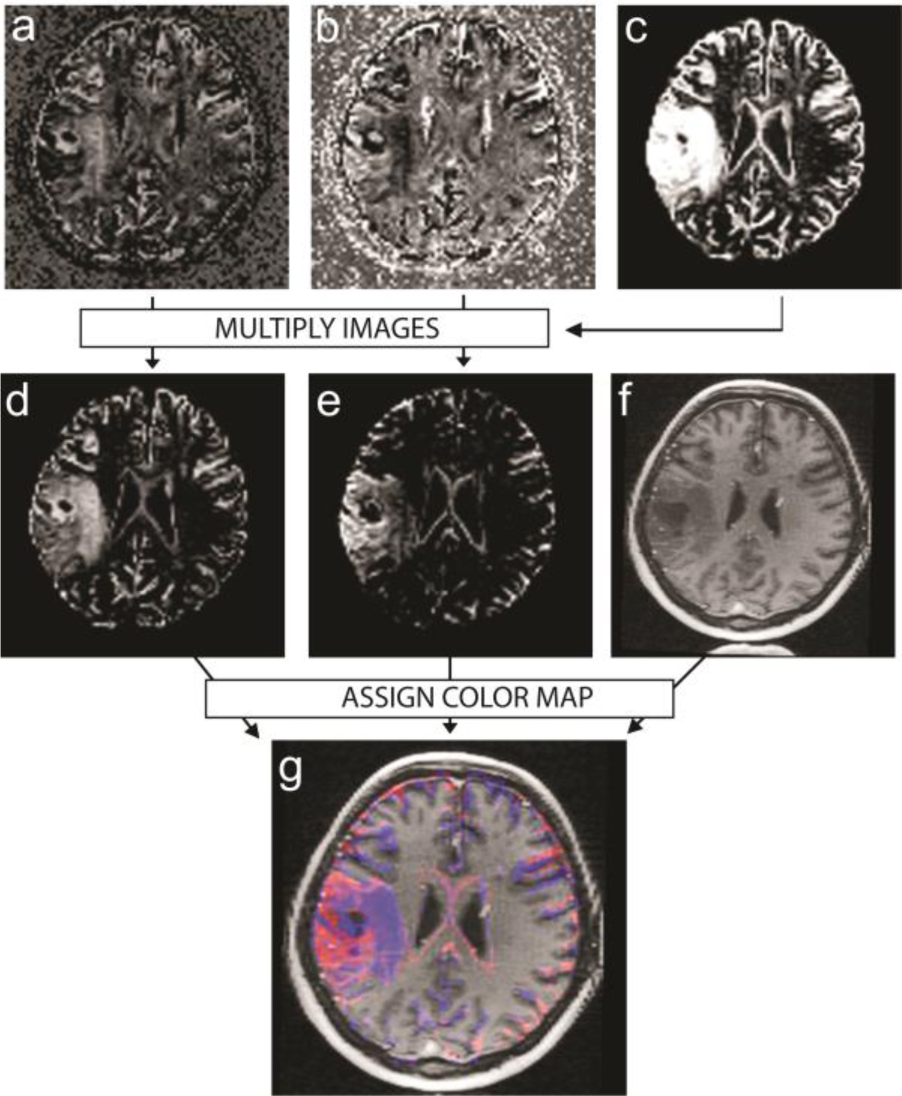
Calculation of intermediate grayscale images and the color-coded glioma grade map. (a) low grade tumor probability map (LGPM), (b) high grade tumor probability map (HGPM), (c) tumor probability map (TPM), (d,e) LGPM and HGPM weighted with the TPM, (f) T1-weighted anatomical image. (g) glioma grade maps are generated by assigning color-code to the probability maps (d,e) and merging them with the postcontrast T1 images.

### Validation of the results

We found it practical to compute a variable that quantifies the overall estimated grade for each tumor volume; this was specified by the values measured over the tumors on the high-grade tumor probability maps, designated as the “grade index”. Due to the fact that the ANN algorithm used a decision value of 0.5 for discriminating the two grades, values from 0 to 0.5 represented low-grade while the 0.5 to 1 range indicated high-grade tumors – this partitioning was also tested by matching with the corresponding histopathology. Glioma grade maps were visually inspected; tumors showing pronounced regional heterogeneity on grade maps were further analyzed by comparing the imaging biomarker values with the microscopic and immunohistochemical findings such as the Mib-1 labeling index.

## RESULTS

Databases were successfully generated by translating voxel-wise imaging and *a priori* grade information into a 2D matrix: database “A” consisted of 162 609 whereas database “B” initially comprised approximately 12 million samples. To reduce the computational burden when working with database “B”, samples were randomly omitted, keeping only 5% of the extratumoral and intracerebral voxels in the database.

The first neural network predicted the grade of voxels inside the tumor borders with 82.12 ± 1.84% accuracy (average of 10 runs, tested on the independent holdout sample, putatively marking the accuracy for new observations). Next, the intra-tumoral voxel membership was estimated correctly in 86.44 ± 0.41% of the samples. Grade index was calculated for each outlined tumor volume. For low grade cases it was 0.281 ± 0.164 (range 0.012 – 0.601) while in high grade lesions it was 0.646 ± 0.148 (range 0.331 – 0.837), the difference was significant (p<0.001, Mann-Whitney U test). Additionally, the grade index showed high correlation with the WHO grade (i.e. II, III or IV); Pearson score: 0.709, p<0.001. With the cut-off point set to 0.5, the grade index could identify high grade cases with 92.31% specificity, 85.71% sensitivity, AUC: 0.967.

Visual assessment of the TPM images, T1 anatomical scans and tumor outline ground truth data revealed good correspondence with the predicted borders, with the following exceptions. Normal-appearing brain regions contained false positive voxels with either blue or red appearance, mainly matching the borders of the gray matter and the cerebrospinal fluid; this error was reported in 8 cases and could presumably be attributed to partial volume or co-registration artifacts (Figure 4/e, white arrow). Six illustrative images of various glioma subtypes and WHO grades were selected to demonstrate the diagnostic features of grade maps (Figure 4).

**Figure 4.**
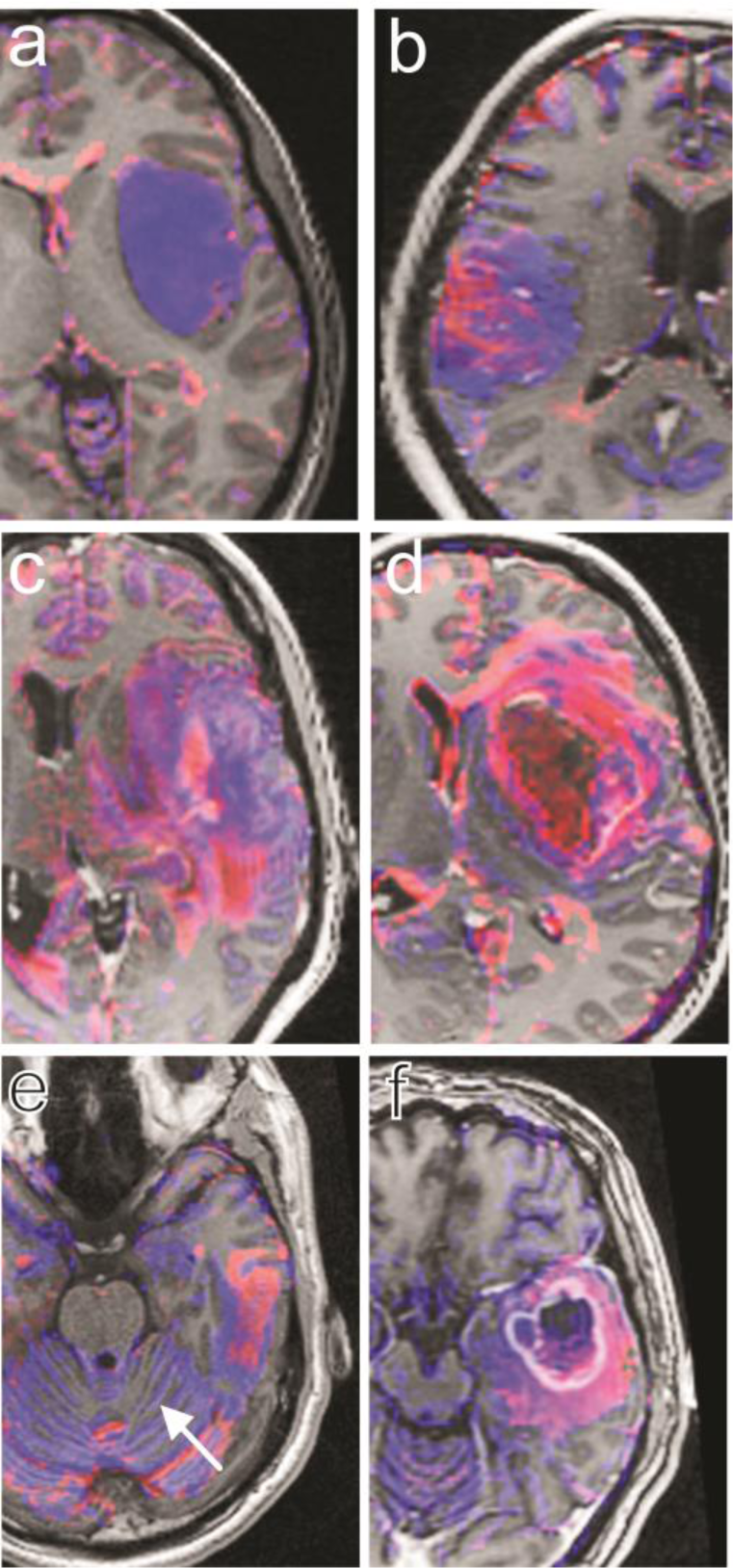
Color-coded glioma grade maps depicting various glioma cases. (a) astrocytoma gr. II. tumors are shown as predominantly blue lesions. (b) oligoastrocytoma gr. II. In a number of cases where the histopathological evaluation judged the lesion as low-grade, the grade maps revealed focal heterogeneity. (c) this astrocytoma gr. III. displays pronounced regional heterogeneity on the glioma grade map; whereas the contrast enhancing regions are well co-localized with the red regions resembling high grade characteristics. (d) glioblastoma multiforme tumor. (e) a misclassified low-grade case with high cellular atypia. Coregistration and partial volume errors are observed outside the lesion (arrow). (f) glioblastoma multiforme with voluminous necrotic areas, incorrectly classified as low-grade.

The appearance of astrocytoma, oligoastrocytoma grade II and oligodendroglioma grade II tumors on the color-coded grade maps was blue (Figure 4/a). Sparse high-grade regions were identified in about six of the 17 non-enhancing and otherwise homogeneous low-grade tumors (Figure 4/b) while the focal heterogeneity as marked by contrast-enhancement was revealed correctly in 77.8% (7/9) by regions of red hue in low grade gliomas. WHO grade III (high grade) oligoastrocytomas (Fig. 4c) and glioblastoma multiformes predominantly appeared purple to red, with marked heterogeneity as indicated by blue patches (Figure 4/d). In 4 of 40 cases, classification by the grade index proved incorrect, for which the following facts are assumed to be responsible. In a patient with a voluminous glioblastoma multiforme (Figure 4/f) this could be putatively ascribed to the relatively high presence of necrotic areas in the tumor, unmasked during the classifier training, hence areas inside necrotic masses were predominantly recognized as low-grade with markedly high-grade rims that closely resembled the contrast-enhancing areas on T1 scans.

In the other misclassified high-grade case, we found no justification for the result; although the designated grade index was just below the cut-off point. The grade indices for the misclassified glioblastoma multiforme tumors were 0.331 and 0.48, respectively. Two low-grade tumors were improperly classified. In one case the pathologist described high Mib-1 labeling index (20%), hyperchromatic nuclei, geometric neovascularisation and a cellular atypia almost reaching the criteria for grade III classification; further on, closer clinical inspection was suggested for the neuro-oncology team. A cross-section image from the grade map of this case is shown in Fig. 4e.

The heterogeneous appearance of many tumors prompted us to investigate histopathological results in detail. It was found that the grade index was significantly higher in the group of tumor samples displaying pathologic endothelial proliferation patterns (p=0.01, Mann-Whitney U test). The grade index showed strong, positive correlation with the categories of Mib-1 labeling index; i.e. low (0-4%), medium (4-10%) or high (>10%). Detailed evaluation of the relationship between tumor characteristics and the grade index is summarized in Table 3.

**Table 3.** The grade index as a quantitative imaging biomarker of gliomas: correlations with histopathology

| Evaluated histopathological characteristics of gliomas | n | Grade index | Level of significance |
| --- | --- | --- | --- |
| WHO grade |  |  |  |
| II. | 26 | 0.281 ± 0.164<br>(0.012 – 0.601) | p<0.001<sup>a</sup><br>(Correlation coefficient=0.709) |
| III. | 3 | 0.672 ± 0.162<br>(0.514 – 0.837) |  |
| IV. | 11 | 0.638 ± 0.151<br>(0.331 – 0.835) |  |
| Patterns of endothelial proliferation |  |  |  |
| Not present / normal | 26 | 0.312 ± 0.206<br>(0.012 – 0.837) | p=0.001<sup>b</sup> |
| Pathologic patterns present | 8 | 0.599 ± 0.134<br>(0.331 – 0.805) |  |
| Proliferation rate: MIB-1 labeling index |  |  |  |
| <4% | 15 | 0.286 ± 0.181<br>(0.016 – 0.631) | p=0.002<sup>a</sup><br>(Correlation coefficient=0.561) |
| 4-10% | 6 | 0.362 ± 0.245<br>(0.137 – 0.667) |  |
| >10% | 6 | 0.606 ± 0.142<br>(0.413 – 0.837) |  |
| p53 positivity |  |  |  |
| <5% | 9 | 0.434 ± 0.22<br>(0.137 – 0.667) | p=0.758<sup>b</sup> |
| >5% | 7 | 0.383 ± 0.177<br>(0.151 – 0.645) |  |

a: Evaluated using the Pearson test of correlation
b: Mann-Whitney-Wilcoxon test

## DISCUSSION

The current gold standard for determining glioma subtype and grade is surgical biopsy, which is subject to sampling errors. Small volume surgical samples may not represent the entire tumor, and due to the marked focal heterogeneity of gliomas, it may lead to the false determination of subtype. MRI guidance addresses this problem by specifying areas which presumably present more malign biological behavior: it is accepted that contrast enhancing regions of gliomas are brought about by the disruption of the blood-brain barrier. Further valuable radiological features of high-grade gliomas on gadolinium-enhanced MR images are signal intensity heterogeneity, necrosis, hemorrhage, degree of oedema and mass effect. To precisely characterize an entity of pronounced heterogeneity like gliomas, further information is required: perfusion studies, MR spectroscopy and diffusion measurements were found feasible for this objective. Such modalities allow calculating functional maps and imaging biomarkers (Galban et al., 2009; Ellingson et al., 2010) and have been shown to play a complementary role in assessing therapy response or pattern of recurrence. (Mardor et al. 2004; Verma et al., 2008; Pope et al., 2009).

In our study, two databases comprising voxel-wise image data of 40 glioma patients were construed, whereas artificial neural network computing was utilized to reclassify the original image voxels and by the same token, it becomes possible to classify voxels of undiagnosed cases. Grayscale images were generated that depicted the probabilities of tumor classification (LGPM and HGPM); and eventually, they were combined to produce color-coded composite images, the grade map (Figure 3). In this image type, purple to red hue suggests higher grade, while low grade tumors mainly appeared blue. This graphical representation may allow fast tumor characterization. The neural network approach was effective in determining tumor grade of individual voxels, whereas a new variable calculated from the voxel-wise outputs of the classifier – the grade index of entire tumor volumes – allowed sufficient classification. In terms of the correct determination of glioma grade, our results exceed the diagnostic power of conventional MR imaging as described by Law et al. (postgadolinium MRI: 72.5% sensitivity, 65% specificity; grade index classification: 85.7% sensitivity, 92.3% specificity); however, it was reported that the feasibility of using perfusion MRI data vastly improves (95% sensitivity; 57.5% specificity). Arvinda and co-authors found that ADC, perfusion measurements and their combination could be successfully employed to characterize glioma grade (Arvinda et al., 2009). Herein we report similar results, the grade index being more specific compared to the ADC values alone (92.3% and 87.1%, respectively). Jakab and co-authors previously described a multivariate discriminant analysis approach that uses histogram descriptors for classifying gliomas, the grade index based classification shows similar sensitivity and higher specificity (Jakab et al., 2011). While conventional MR imaging provides usable features to discriminate grade IV (GBM) tumors from grade II malignancies, the separation of grade III. anaplastic astrocytomas from low grades is inefficient; White and co-authors described that fractional anisotropy (FA) values and descriptors of the distribution of such values over the tumor volume can increase the sensitivity of grade II – III discrimination (White et al., 2011). Our method provides a novel way to incorporate FA as a feature.

Our investigation has several limitations. The reproducibility of the artificial neural network (ANN) algorithm is often disputed; it is generally considered as a “black box” rather than an analytical approach. Increasing the number of processing layers in the ANN will reduce the classification error but consequently causes a loss of generalizability (Hagberg et al., 1998). In our investigation, the number of samples (i.e. voxels) was high and the resulting network structure was kept simple, hence we conclude that the network is not overtrained. It is believed that reproducibility issues would partially be resolved by employing other algorithms such as support vector machines which has already been shown promising in glioma grading (Li et al., 2006). Also, it would be desirable to perform the longitudinal scanning and grade map calculation of a single patient, to evaluate the inter-subject reproducibility of the proposed method. We used a split of training, testing and holdout samples to avoid evaluation on the same samples used for training. Nevertheless, prospective clinical testing is necessary to evaluate whether a radiologist can perform better with the presented tool than without it. We hypothesized that during the training procedure it is feasible to assign the same categorical diagnosis for each voxel in one particular tumor; however, this presumption required that pathological diagnoses were made from the analyses of representative tissue samples. Matching a specific set of voxels to the position of the surgical sampling would enable better correlation of voxel-wise imaging data and tumor grade. If the assumption is true that the grade index is a quantitative biomarker for depicting alterations in glioma microstructure representative for biological progression, it may also be hypothesized that the values of this biomarker for grade III tumors are between the values of grade II and IV gliomas. Albeit this was not confirmed by our study, the two grade III case had higher grade indices compared to low grade samples: 0.673 ± 0.161 and 0.281 ± 0.164. This unusual distribution of grade indices in grade III tumors could be attributed to the low number of cases. Another limitation in our study design is the inclusion of tumors with mixed tissue composition like oligodendrogliomas; it is not evident that the same characteristic changes occur in terms of diffusion or relaxation parameters during the transition from any glioma subtype to higher grades therefore making it harder to generalize this phenomena.

De Edelenyi and colleagues found that multidimensional MRI data could be used to create images demonstrating the classification or “nosology” of brain neoplasms; moreover, they suggested incorporating diffusion data in similar future studies. To the best of our knowledge, this is the first study that performs glioma characterization using machine-learning algorithms that combine imaging data of T1- and T2-weighted, diffusion anisotropy and apparent diffusion coefficient information.

Grade maps are graphical representations of tumor subtype and heterogeneity whilst the grade index was defined as an overall estimate of tumor grade as determined by the assignments of classifiers. In a number of cases, our findings allowed identification of tumors with prominent regional heterogeneity and marked biological progression. The glioma grade map and grade index might serve as imaging biomarkers for the characterization of brain gliomas and complement preoperative information available for clinicians.

